# Keratin 19 maintains E-cadherin localization at the cell surface and stabilizes cell-cell adhesion of MCF7 cells

**DOI:** 10.1101/2020.05.28.119297

**Authors:** Sarah Alsharif, Pooja Sharma, Karina Bursch, Rachel Milliken, Meagan Collins, Van Lam, Arwa Fallatah, Thuc Phan, Priya Dohlman, Georges Nehmetallah, Christopher B. Raub, Byung Min Chung

## Abstract

A cytoskeletal protein keratin 19 (K19) is highly expressed in breast cancer but its effects on breast cancer cell mechanics are unclear. Using *KRT19* knockout (KO) cells and cells where K19 expression was rescued, we found that K19 is required to maintain rounded epithelial-like shape and tight cell-cell adhesion of MCF7 cells. A loss of K19 resulted in a lower level of plakoglobin and internalization of E-cadherin in early and recycling endosomes. Inhibiting internalization restored cell-cell adhesion of *KRT19* KO cells, suggesting E-cadherin internalization contributes to defective adhesion. Ultimately, while K19 inhibited cell migration, it was required for cells to form colonies in suspension. Our results suggest that K19 stabilizes E-cadherin complexes at the cell membrane to maintain cell-cell adhesion which inhibits cell migration but provides growth and survival advantages for circulating tumor cells. These findings provide context-dependent roles of K19 during metastasis.

## Introduction

The majority of cancer deaths are due to metastasis ^1–3^. Tumor metastasis is a multistep process that involves spreading of cancer cells from a primary site to colonization at a distal site ^2,4^. During its initiation, metastasis has been linked closely to epithelial-to-mesenchymal transition (EMT) whereby cells lose morphological traits of epithelial cells, including tight cell-cell adhesion and apical-basal polarity, and gain mesenchymal traits such as increased ability to undergo migration and invasion ^2,5,6^. At the molecular level, epithelial markers such as E-cadherin and keratins become downregulated while mesenchymal markers such as vimentin become upregulated during EMT.

Keratins belong to an intermediate filament family of cytoskeletal proteins, and keratin filaments maintain epithelial cell polarity and mechanical integrity through intercellular and cell-extracellular matrix junctions called desmosomes and hemidesmosomes, respectively ^7,8^. Decreased expression of keratins during EMT is considered to contribute to an initiation of metastasis by loosening cell-cell attachment through disassembly of desmosomes ^9–11^. Therefore, it has been suggested that maintenance of intercellular adhesion by keratins serves as a barrier against EMT and cell migration ^11^ a concept supported by several *in vitro* studies involving keratins expressed in simple epithelium and keratinocytes ^5,12–14^. However, upregulation of select keratins has been shown to enhance cell migration and invasion in certain cancer settings ^12,15,16^, likely due to the fact that some cancer cells invade extracellular matrix collectively ^17,18^.

Following initiation, metastatic cells intravasate into the blood stream and must survive in suspension as circulating tumor cells (CTCs) en route to a distal site ^2,4,19^. High metastatic potential of CTCs has been associated with stem-like properties ^20^ and also with clusters of cells with higher levels of cell adhesion molecules plakoglobin or E-cadherin ^21,22^. However, the role of keratins on stem-like traits and clustering of cancer cells upon detachment from the extracellular matrix remains unclear.

In the context of breast cancer, previous studies have shown that depletion of K19 increases cell migration and invasion *in vitro*, potentially through upregulation of Akt and Notch signaling pathways ^23–25^. However, mammary stem/progenitor cells transformed with sets of oncogenes including mutant Ras and p53 were more metastatic when K19 was present ^26^. In addition, the role of K19 in cell-cell and cell-substrate mechanics contributing to migration of cancer cells has not been defined.

To study the role of K19 on processes fundamental to metastasis, we examined the luminal-subtype MCF7 breast cancer cell line which expresses high levels of K19 ^27,28^. MCF7 cells maintain polarized epithelial phenotype with intact cell-cell adhesions made of desmosomes and adherens junctions ^29–31^. Using MCF7 cells with complete ablation of K19 ^32^ and *KRT19* knockout (KO) cells with re-expression of K19, we observed that K19 is required for the epithelial-like cell shape and proper cell-cell adhesion. These events were accompanied by lower levels of plakoglobin but accumulation of E-cadherin in endocytic compartments in the absence of K19. Importantly, while we confirmed the inhibitory role of K19 on cell migration, K19 was found to be required for cells to grow in low attachment conditions.

## Results

### KRT19 KO cells display an elongated phenotype

Under the microscope, MCF7 *KRT19* KO cells showed a considerable difference in morphology from their parental counterpart. While parental (P) MCF7 cells were mostly epithelial-like and rounded in shape, *KRT19* KO cells exhibited more mesenchymal-like morphology with elongated and spindled shapes (Figure 1A-B). Of note, we examined two *KRT19* KO clones to confirm phenotypes associated with a loss of K19. To quantify the difference in shapes between parental and *KRT19* KO cells, we sorted cells into two categories, rounded and elongated, based on their shapes (Figure 1C & S1A). Scoring of cell shapes confirmed that *KRT19* KO cells were more elongated than parental cells (Figure 1D). Of note, while two *KRT19* KO clones exhibited subtle differences from each other, both were more elongated than parental cells. We then used digital holographic microscopy (DHM) to quantitate morphologies of parental and *KRT19* KO cells (Figure 1E). DHM measured cells based on index of refraction and physical thickness ^33,34^ and gave 17 physical properties at a single cell level (Table S1). Optical phase measurements included pixel mean, standard deviation, and texture parameters, while geometric parameters included cell circularity, eccentricity and form factor. While cell area and perimeter were larger in *KRT19* KO cells, a higher eccentricity and a lower form factor confirmed the elongated phenotype of *KRT19* KO cells (Figure 1F-I).

**Figure 1.**
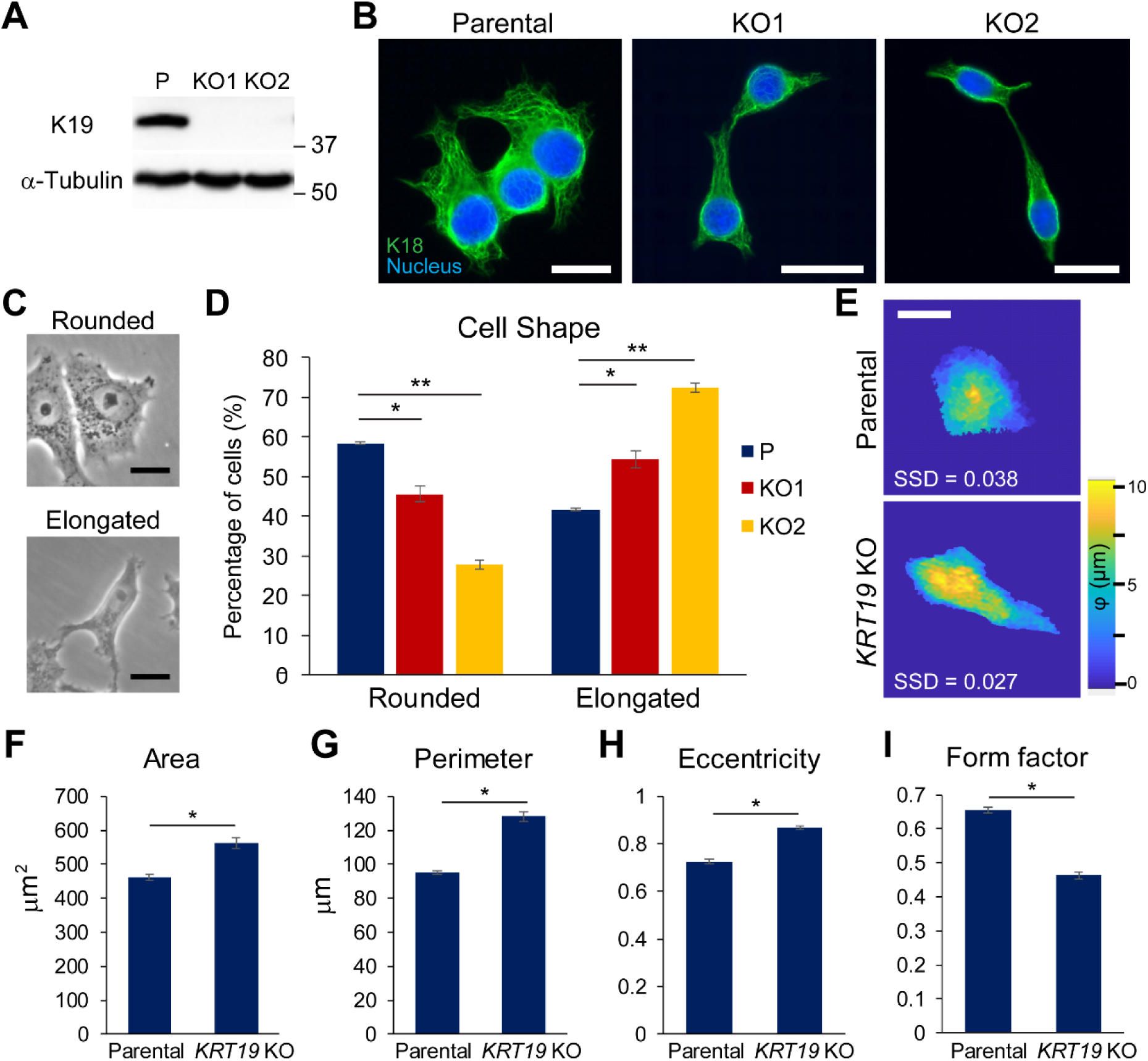
Keratin 19 knockout cells display an elongated phenotype. (**A**) Whole cell lysates of parental (P) control and two different clones (KO1 and KO2) of *KRT19* KO cell lines were harvested, and immunoblotting was performed with antibodies against the indicated proteins. (**B**) Immunostaining of K18 (green) in P and *KRT19* KO cells. Nuclei are shown in blue. Bar, 20 µm. (**C**) Phase contrast images of representative rounded and elongated shapes. Bar, 20 µm. (**D**) Percentages of P and *KRT19* KO cells with rounded or elongated cell shape. Data from five experimental repeats are shown as mean ± SEM. Differences are not statistically significant unless denoted by *p < 0.05; **p < 0.001. (**E**) Phase height maps of P and *KRT19* KO (KO2) cells collected by digital holographic microscopy (DHM). Representative P and *KRT19* KO cells with the smallest sum of squared deviations (SSD) of 17 phase parameters collected shown. A color bar indicates cell phase height. Bar, 10 µm. (**F**) Area, (**G**) Perimeter, (**H**) Eccentricity, and (**I**) Form factor of P and *KRT19* KO cells from DHM analyses. Form factor was calculated using the mathematical formula 4π(area)/(perimeter)^2^. Form factor of 1 represents a perfect circle. Data from three experimental replicates are shown as mean ± SEM. Differences are not statistically significant unless denoted by *p < 0.05

### Weakened cell-cell adhesions in KRT19 KO cells

When culturing cells, we noticed that elongated *KRT19* KO cells were forming loose contacts between neighboring cells whereas parental MCF7 cells were in close contacts with their neighbors (Figure 1B). To quantitate the difference, cell-cell adhesion made by subconfluent parental and *KRT19* KO cells were examined. Adhesions were categorized into three different degrees: high indicates a cell attached to its neighboring cells by making contiguous contacts all along adjoining sides; low indicates a cell attached to its neighbor only by pointed contacts; and medium as those harboring both high and low adhesions (Figure 2A). In the absence of K19, a lower percentage of cells exhibited high adhesion, but more cells showed low adhesion, confirming the observation that a loss of K19 resulted in decreased cell-cell adhesion (Figure 2B).

**Figure 2.**
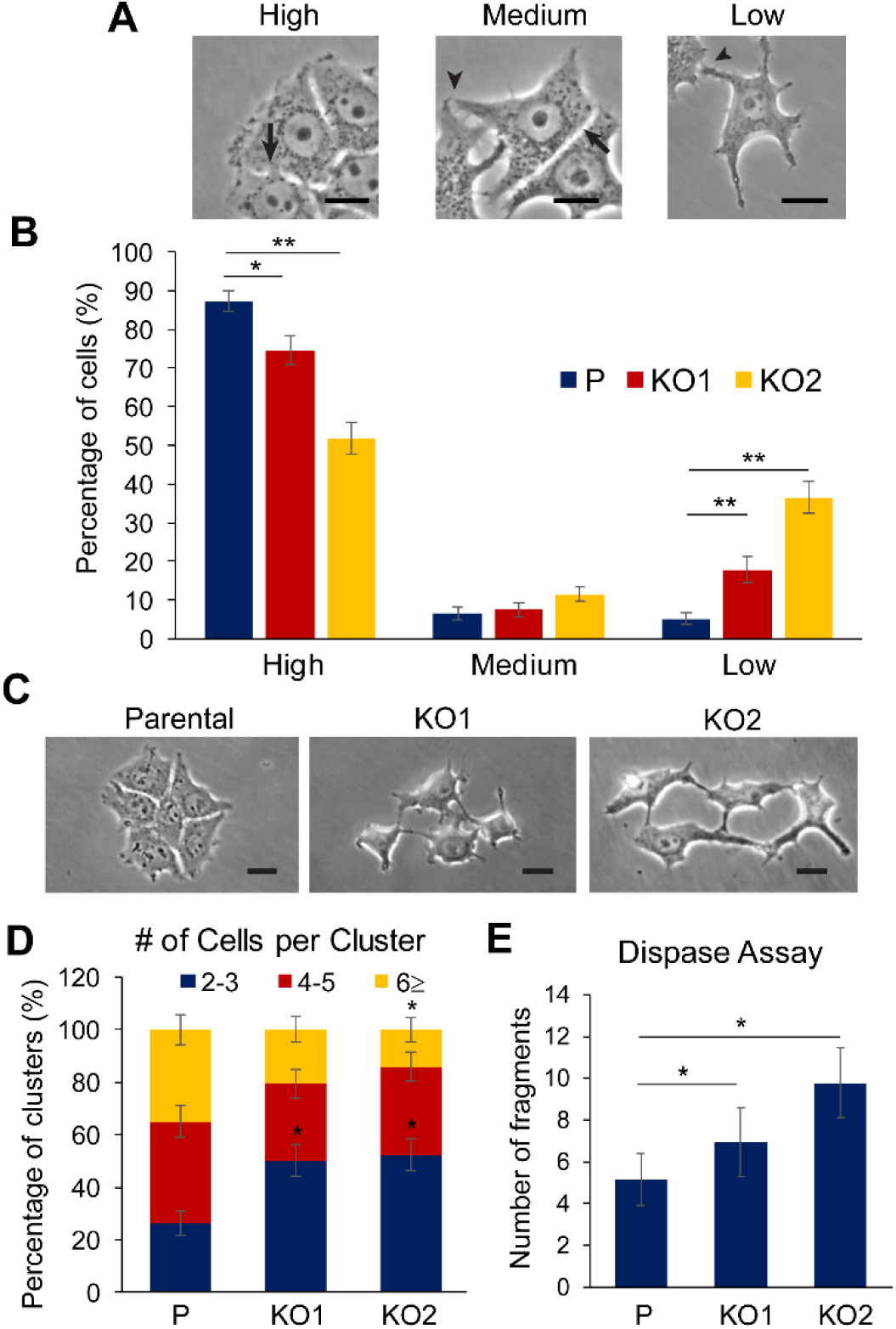
Weakened cell-cell adhesion in *KRT19* KO cells. (**A**) Phase contrast images of representative cells engaged in high, medium and low adhesions. Bar, 20 µm. (**B**) Percentage of parental (P) control and *KRT19* KO clones in high, medium or low adhesions. Data from five experimental repeats are shown as mean ± SEM. Differences are not statistically significant unless denoted by *p < 0.05; **p < 1 × 0.001. (**C**) Phase contrast images of cell clusters. Bar, 20 µm. (**D**) Number of cells per cluster. Clusters classified into three different ranges: 2-3 cells/cluster; 4-5 cells/cluster; and >6 cells/cluster. Data from three experimental repeats are shown as mean ± SEM. Any statistically significant difference between P and *KRT19* KO clone is denoted by * (p < 0.05) above the cluster of *KRT19* KO clone. (**E**) Number of monolayer fragments formed from the dispase assay. Data from at least three experimental repeats, each with three replicates, are shown as mean ± SEM. Differences are not statistically significant unless denoted by *p < 0.05

Since parental MCF7 cells grow in tight clusters even at low confluency, cell adhesion was also assessed by counting number of cells in each cluster after passaging (Figure 2C). The number of cells making contacts as a group was fewer in *KRT19* KO cells compared to parental cells (Figure 2D). In addition, disrupting cell-cell adhesion with dispase treatment and mechanical force induction showed that K19 was required for proper cell-cell adhesion as indicated by a higher number of fragmented monolayers in *KRT19* KO cells (Figure 2E & S1). Collectively, these data support the notion that K19 is required for proper adhesion between cells.

### Decreased cell surface localization of E-cadherin in KRT19 KO cells

In our previous work, we had performed RNA-sequencing to identify genes differentially expressed between parental and *KRT19* KO cells ^32^. Consistent with decreased cell-cell adhesion, *KRT19* KO cells expressed reduced levels of plakoglobin, a desmosomal and adherens junction component (Figure 3A-B) ^32^. Surprisingly however, a functional enrichment analysis using Database for Annotation, Visualization and Integrated Discovery (DAVID) showed that many genes upregulated in *KRT19* KO cells were associated with functions in cell membranes, cell adhesions, and cell junctions (Figure S2). Indeed, although levels of β-catenin, a K19-interacting component of adherens junction (Figure S3) ^24^, remained the same, we found higher levels of E-cadherin, a Ca^2+^-dependent adhesive molecule and a key regulator of cell morphology ^35^, in *KRT19* KO cells (Figure 3A-B). However, as assessed by immunostaining of cells (Figure 3C) and surface biotinylation (Figure 3D-E), the cell surface localization of E-cadherins was decreased in *KRT19* KO cells. Immunostaining also showed that cell surface localization of other cell junctional proteins including plakoglobin, β-catenin and p120-catenin were decreased in *KRT19* KO cells as well (Figure S4). Furthermore, interaction between E-cadherin and β-catenin was lower in *KRT19* KO cells, suggesting a requirement for K19 in proper maintenance of adherens junction (Figure 3F-G). We then decided to test functions of Ca^2+^-dependent adhesion molecules in parental and *KRT19* KO cells. While stimulating Ca^2+^-deprived parental cells with Ca^2+^ allowed cells to re-adhere to each other, Ca^2+^ stimulation had little effect on adherence of *KRT19* KO cells (Figure 3H). These observations suggest that K19 is required for the formation and function of calcium-dependent junctional complexes. Defective cell-cell adhesion in *KRT19* KO cells was also observed upon serum stimulation (Figure S5). Altogether, these data suggest that adherens junctions are not properly maintained in *KRT19* KO cells

**Figure 3.**
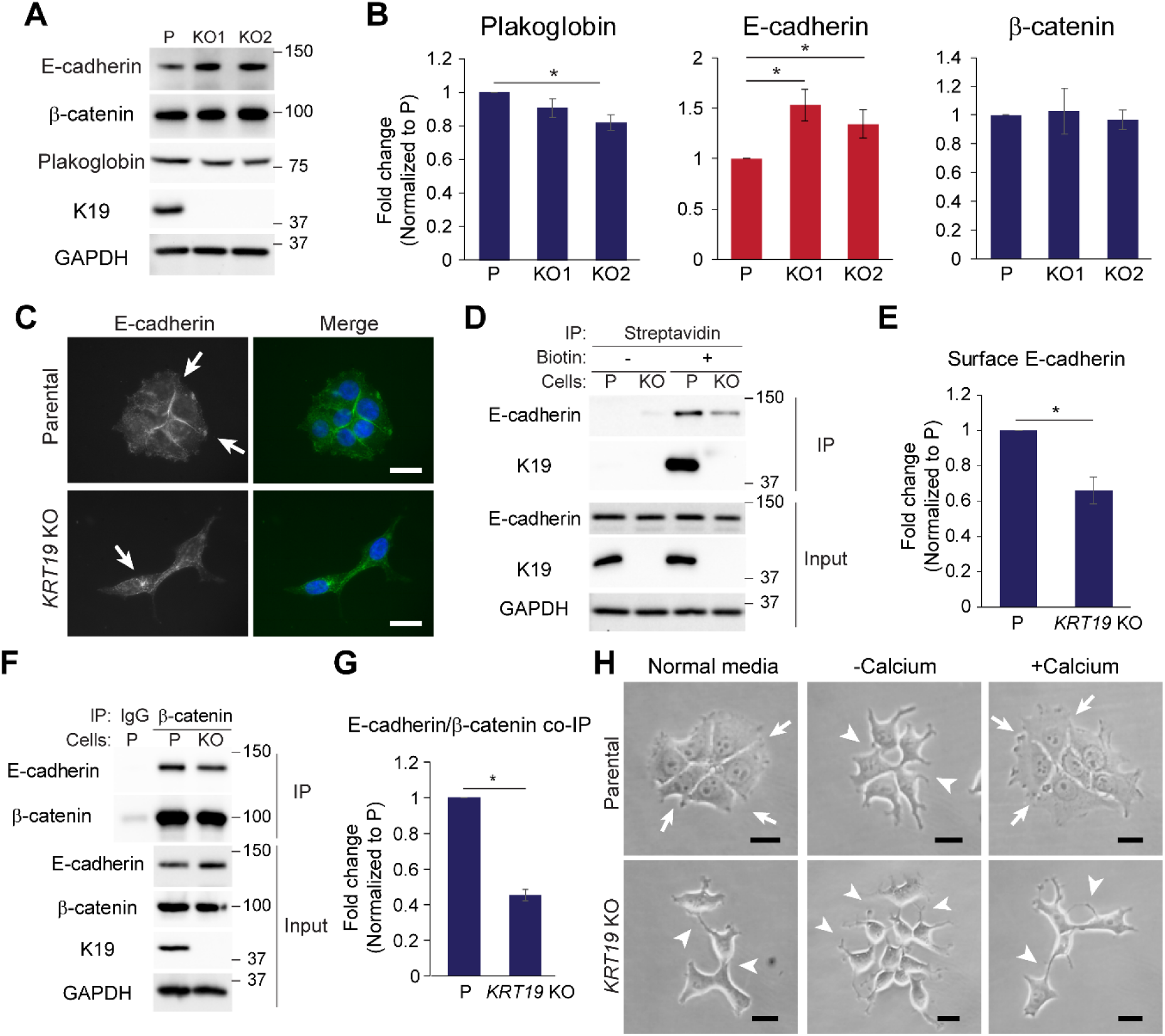
Defective regulation of E-cadherin in *KRT19 K*O cells. (**A**) Whole cell lysates of parental (P) control and *KRT19* KO clones were prepared and immunoblotting was performed with indicated antibodies. Molecular weight in kDa (**B**) Signal intensities of Plakoglobin, E-cadherin and β-catenin from (**A**) were quantified and normalized to those of the GAPDH loading control. Data from at least three experimental repeats normalized to that of the parental control are shown as mean ± SEM. Differences are not statistically significant unless denoted by *p < 0.05. (**C**) Immunostaining of E-cadherin (green) in P and *KRT19* KO (KO2) cells. Nuclei are shown in blue. Arrows indicate E-cadherin localization. Bar, 20 µm. (**D**) Streptavidin beads were used to pulldown P and *KRT19* KO (KO2) cells either treated (+) or untreated (-) with biotin for cell surface labelling of proteins. Immunoprecipitates (IP) and inputs were subjected to SDS-PAGE and immunoblotting was performed with antibodies against the indicated proteins. (**E**) Signal intensities of E-cadherin in IP of biotin-treated samples from (**D**) were quantified and normalized to those of E-cadherin in input. Data from at least three experimental repeats normalized to that of the parental control are shown as mean ± SEM. *p < 0.05. (**F**) Co-IP of β-catenin with E-cadherin. IP with anti-β-catenin antibody or IgG control was perform in P and *KRT19* KO (KO2) cells. Immunoprecipitates (IP) and inputs were subjected to SDS-PAGE and immunoblotting was performed with antibodies against the indicated proteins. (**G**) Signal intensities of E-cadherin in IP from (**F**) were quantified and normalized to those of β-catenin in IP. Data from at least three experimental repeats normalized to that of the parental control are shown as mean ± SEM. *p < 0.05. (**H**) Phase contrast images of P and *KRT19* KO (KO2) cells. Cells were either grown in normal growth condition (Normal media), placed in calcium-free media for 6 h, then left unstimulated (-Calcium) or stimulated (+Calcium) with CaCl_2_ for 4 h. Arrows indicate high cell-cell adhesions and arrowheads indicate low cell-cell adhesions. Bar, 20 µm.

### Internalization of E-cadherin into endocytic compartments in KRT19 KO cells

E-cadherin constantly undergoes endocytosis and recycling in the absence of stable cell-cell contacts ^36^. The fact that total E-cadherin level is higher while its surface level is lower in *KRT19* KO cells indicates that most E-cadherin is localized in endocytic compartments. In order to determine the exact location of E-cadherin in *KRT19* KO cells, we performed co-immunostaining of E-cadherin and endocytic markers. Cells were stimulated with labelled transferrin to mark early/recycling endosomes (Figure 4A) while LAMP1 was used as a marker of late endosomes/lysosomes (Figure 4B). Results show that while E-cadherin colocalized with labelled transferrin in *KRT19* KO cells, very little, if any, E-cadherin colocalized with LAMP1, suggesting that E-cadherin in *KRT19* KO cells was localized to early/recycling endosomes. Indeed, when *KRT19* KO cells were treated with a dynamin inhibitor dynasore to inhibit E-cadherin internalization ^37^, *KRT19* KO cells showed marked improvement in cell-cell adhesion (Figure 4C). These data suggest that E-cadherin is internalized to early/recycling endosomes in the absence of K19 (Figure 4D).

**Figure 4.**
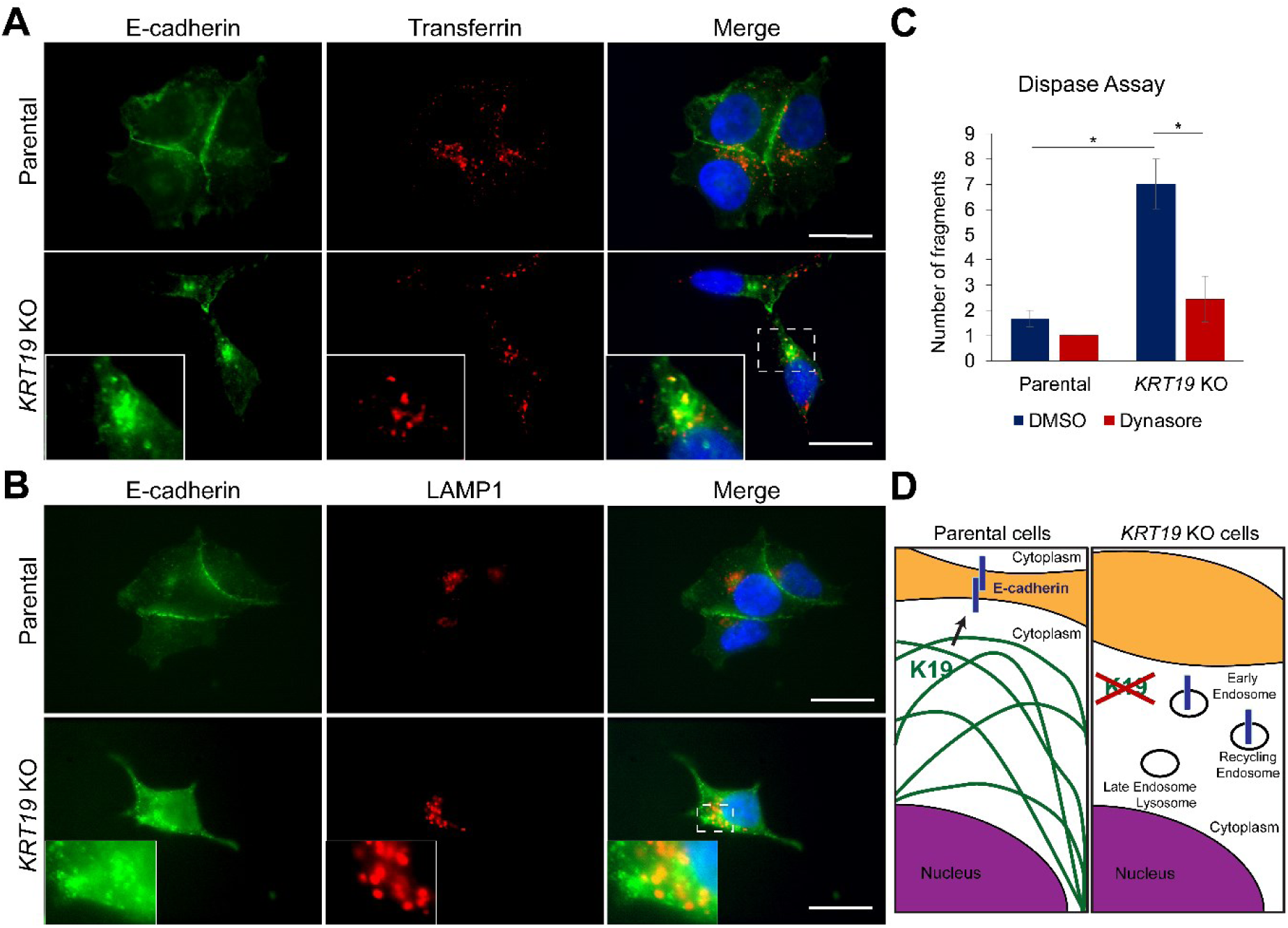
Inhibiting internalization rescues cell-cell adhesion of *KRT19* KO cells. Parental and *KRT19* KO (KO2) cells were either (**A**) incubated with labelled transferrin (red) for 30 min, then immunostained for E-cadherin (green) or (**B**) co-immunostained for E-cadherin (green) and LAMP1 (red). Nuclei are shown in blue. Insets: areas in *KRT19* KO cells rich in E-cadherin localization. Bar, 20 µm. (**C**) Cells treated with dynamin inhibitor (Dynasore) or DMSO control for 2 h were subjected to dispase assay. Data from at least three experimental repeats, each with three replicates, are shown as mean ± SEM. Differences are not statistically significant unless denoted by *p < 0.05. (**D**) Schematic of how K19 influences E-cadherin localization. While E-cadherin is localized to cell surface in the presence of K19 (left panel), it is internalized and accumulated in early/recycling endosomes in the absence of K19 (right panel).

### K19 re-expression rescues defects in in KRT19 KO cells

To confirm that altered morphology (Figure 1) and cell-cell adhesion (Figure 2) in *KRT19* KO cells were due specifically to the absence of K19, we decided to examine cells upon re-expression of K19. Introducing GFP-tagged K19 in *KRT19* KO cells reverted cells into rounded shape (Figure 5A). Quantitation of cell shape, as was done in Fig 1C, confirmed significant increase in cells with rounded shape upon expression of GFP-K19 in *KRT19* KO cells compared to the GFP control (Figure 5B). Furthermore, *KRT19* KO cells stably expressing K19 were more resistant to dispase-induce fragmentation compared to the vector control (Figure 5C), confirming the role of K19 in maintaining cell-cell adhesion. Re-expression of K19 also raised levels of plakoglobin while lowering levels of E-cadherin (Figure 5D), consistent with altered levels of plakoglobin and E-cadherin observed in *KRT19* KO cells (Figure 3A-B). Finally, expression of GFP-K19 in *KRT19* KO cells increased E-cadherin co-immunoprecipitating with β-catenin, suggesting that re-expression of K19 increased formation of the adherens junction complex (Figure 5F). Altogether, these data confirm that K19 is required for proper cell shape and cell-cell adhesion while maintaining plakoglobin level and E-cadherin-β-catenin complex.

**Figure 5.**
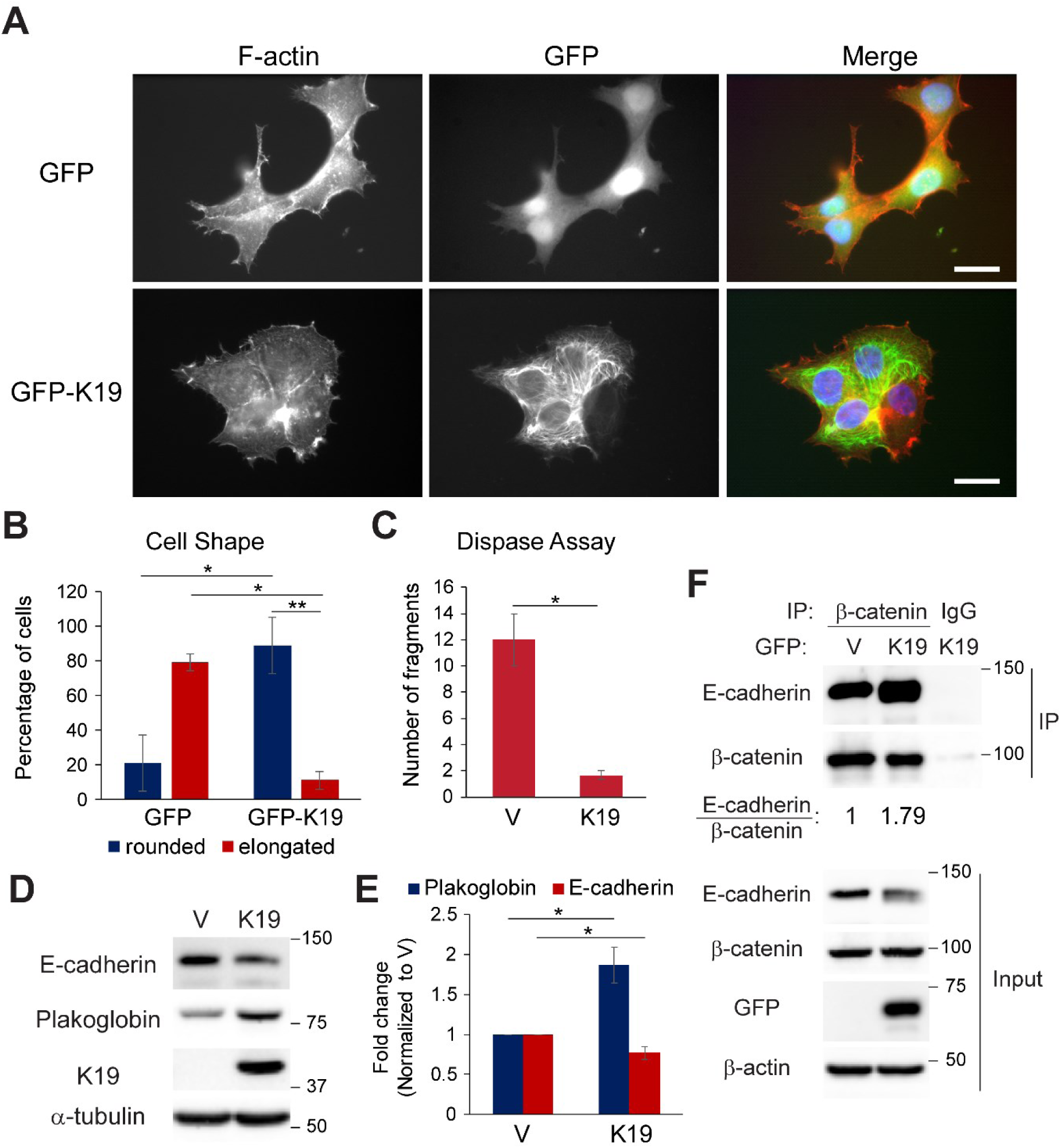
K19 re-expression rescues defects associated with *KRT19* KO cells. (**A**) *KRT19* KO (KO2) cells stably expressing GFP or GFP-K19 were co-immunostained for GFP (green) and F-actin (red). Nuclei are shown in blue. Bar, 20 µm. (**B**) Percentage of cells from (**A**) with rounded or elongated cell shape. Data from five experimental repeats are shown as mean ± SEM. Differences are not statistically significant unless denoted by *p < 0.05; **p < 1 × 0.001. (**C**) Number of monolayer fragments formed by *KRT19* KO (KO2) cells stably expressing vector control (V) or K19 from the dispase assay. Data from at least three experimental repeats, each with three replicates, are shown as mean ± SEM. *p < 0.05. (**D**) Whole cell lysates of *KRT19* KO cells stably expressing vector control (V) or K19 were prepared and immunoblotting was performed with indicated antibodies. (**E**) Signal intensities of Plakoglobin and E-cadherin from (**D**) were quantified and normalized to those of the GAPDH loading control. Data from at least three experimental repeats normalized to that of the vector control are shown as mean ± SEM. *p < 0.05. (**F**) Co-IP of β-catenin with E-cadherin in *KRT19* KO cells stably expressing GFP or GFP-K19. IP with anti-β-catenin antibody or IgG control was perform. Immunoprecipitates (IP) and inputs were subjected to SDS-PAGE and immunoblotting was performed with antibodies against the indicated proteins. Signal intensities of E-cadherin in IP from were quantified and normalized to those of β-catenin in IP and GFP control.

### K19 inhibits cell migration but is required for the anchorage-independent growth

Cell morphology and adhesion of cancer cells are linked to events during metastasis such as cell migration, survival and growth in low adherence conditions. Therefore, we decided to examine how K19 affects these cancer cell behaviors. In wound closure assays, *KRT19* KO cells closed scrape wounds faster compared to parental cells (Figure 6A-B) while re-expression of K19 in *KRT19* KO cells slowed wound closure (Figure 6C), confirming the inhibitory role of K19 in cell migration. Transwell migration assays also confirmed the inhibitory role of K19 in cell migration, as migration was enhanced in *KRT19* KO cells but was suppressed upon expression of GFP-K19 (Figure 6D-E). Interestingly however, when cells were placed on low attachment plates to assess growth in suspension ^38^, mammosphere formation was compromised in *KRT19* KO cells (Figure 7A-B). Re-expression of K19 using GFP-K19 confirmed that K19 is required for mammosphere formation on low attachment plate (Figure 7C-D). Similarly, growing cells in soft agar also showed that K19 is responsible for the formation of colonies in anchorage-independent conditions as colony area was less in *KRT19* KO cells (Figure 7E-F) and reduced formation of colonies was rescued upon GFP-K19 re-expression in *KRT19* KO cells (Figure 7G). Taken together, these data demonstrate that K19 hinders cell migration but promotes colony formation in suspension.

**Figure 6.**
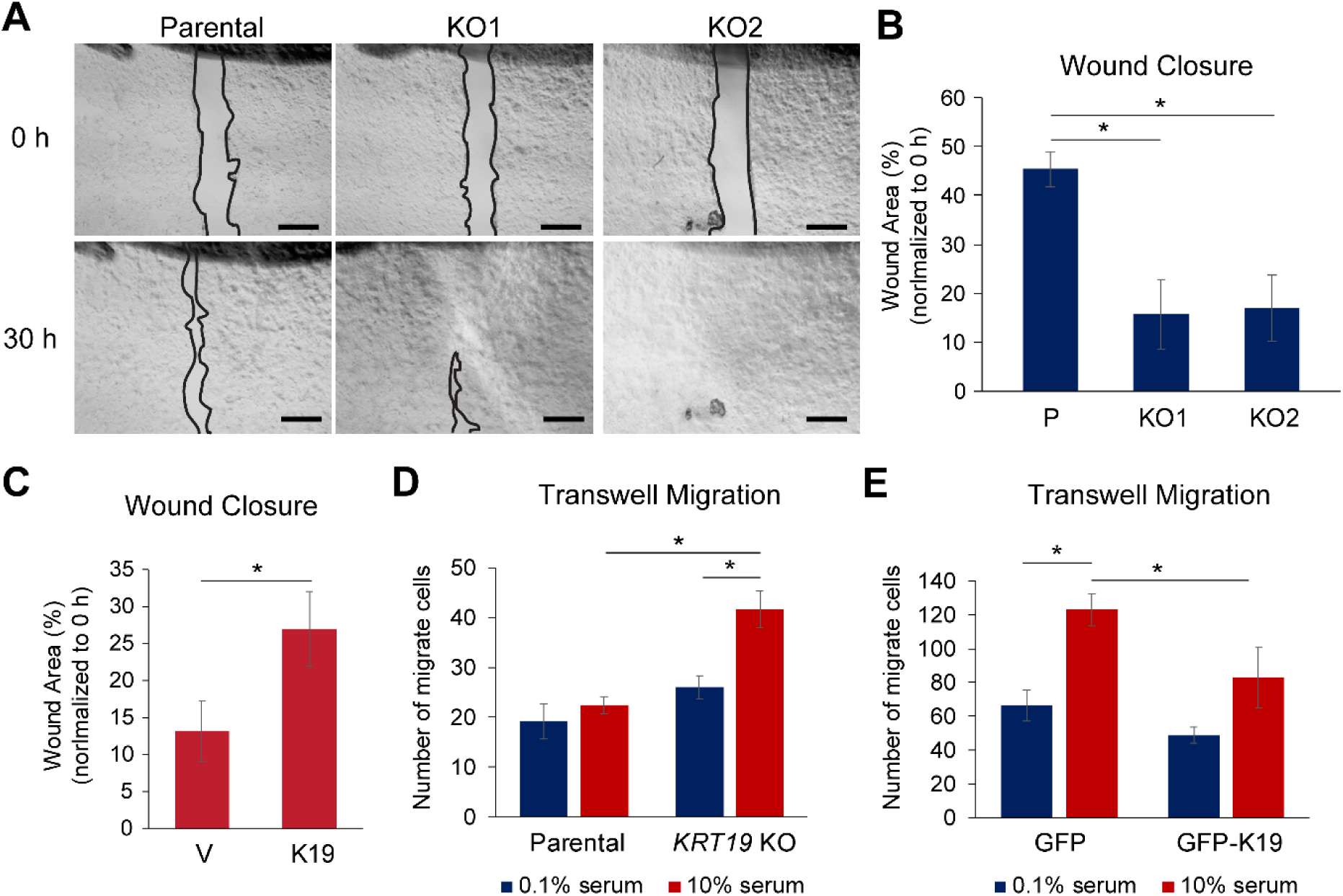
K19 inhibits cell migration. (**A**) Wound closure of parental control and *KRT19* KO clones. Phase contrast images of wound area at 0 and 30 h after scratch. Bar, 0.5 mm. (**B**) Wound areas from (**A**) were quantitated using the ImageJ software and normalized to that at 0 h. (**C**) Wound closure of *KRT19* KO (KO2) cells stably expressing vector control (V) or K19. Wound areas were quantitated using the ImageJ software and normalized to that at 0 h. Transwell migration of (**D**) parental and *KRT19* KO (KO2) cells or (**E**) *KRT19* KO (KO2) cells stably expressing GFP or GFP-K19 in the presence of either 0.1 or 10 % serum as chemoattractant. Migrated cells were identified either by staining nuclei with propidium iodide or using GFP signals under a fluorescent microscope. Number of migrated cells per high-power field were quantified the ImageJ software. For all assays, data from three experimental repeats, each with three replicates, are shown as mean ± SEM. * p < 0.05

**Figure 7.**
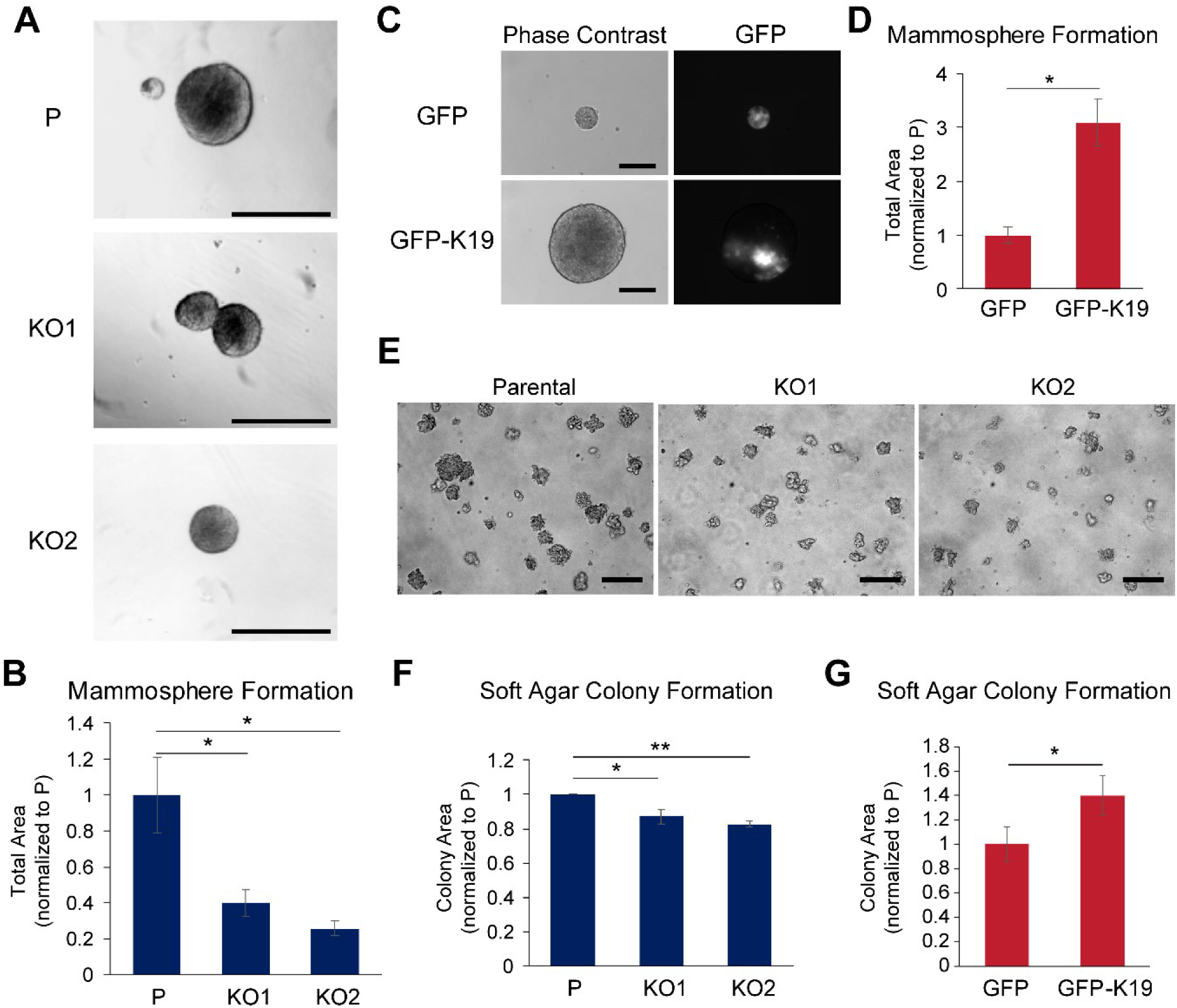
K19 is required for the anchorage-independent growth of MCF7 cells. (**A**) Mammosphere formation of parental (P) control and *KRT19* KO clones. Phase contrast images of mammospheres grown in ultra-low attachment plates for 7 days. Bar, 100 µm. (**B**) Mammosphere areas from (**A**) were quantitated using the ImageJ software. (**C**) Phase contrast and immunofluorescence images of mammospheres formed by *KRT19* KO (KO2) cells stably expressing GFP or GFP-K19 in ultra-low attachment plates. Bar, 100 µm. (**D**) Mammosphere areas from (**C**) were quantitated using the ImageJ software. (**E**) Anchorage-independent growth of parental and *KRT19* KO clones. Phase contrast images of colonies grown in soft agar for 7 days. Bar, 200 µm. (**F**) Colony areas from using (**E**) were quantitated using the ImageJ software. (**G**) *KRT19* KO cells stably expressing GFP or GFP-K19 in soft agar for 7 days and colony areas were measured using the ImageJ software. For all assays, data from three experimental repeats, each with three replicates, are shown as mean ± SEM. * p < 0.05; **p < 0.001

## Discussion

In this study, we demonstrate the role of K19 in maintaining rounded epithelial cell morphology of the MCF7 breast cancer cell line, which is of estrogen-positive luminal subtype where predominant expression of K19 can be found ^39–43^. Interestingly, altered cell shape due to changed K19 expression was also observed in transformed mammary stem/progenitor cells ^44^ and triple negative breast cancer cell lines BT549 ^23^ and MDA-MB-231 ^45^, suggesting that K19 plays a crucial role in maintaining the architecture of breast cancer cells across different differentiation stages and molecular subtypes. Although functions of K19 in normal settings have not been fully resolved, K19 is likely involved in maintaining cell architecture through desmosomes as a member of the keratin family of proteins. Still, there are other keratins present in cells to compensate for the loss of K19, and a loss of all keratins did not by itself alter the shape of tumor cells in lung ^46^. Therefore, it is rather intriguing that the absence of K19 can have such profound effects on breast cancer cell morphology.

Along with decreased cell-cell adhesion, MCF7 cells lacking K19 showed more migration, suggesting that the maintenance of cell-cell adhesion by K19 helps cancer cells retain epithelial morphology and inhibits cancer cell migration from the primary tumor site. Such function would be consistent with the well-known role of keratins during EMT. Of note, vimentin overexpression also caused MCF7 cells to become elongated and more motile ^47^. Therefore, K19 may be an integral member of keratins whose levels relative to vimentin govern the morphology and motility of epithelial cells.

Despite faster migration, *KRT19* KO cells were less efficient in forming mammospheres on low attachment plates and colonies in soft agar, conditions where cells were subjected to growth without being able to adhere to extracellular matrix. Anchorage-independent cell growth has been shown to correlate with metastatic potential as it mimics conditions that circulating tumor cells (CTCs) encounter inside the vasculature ^48^. Our findings suggest that strong cell-cell adhesion made by K19 provides survival and growth advantages to cancer cells, thus increases metastatic potential of CTCs. Indeed, it has been shown that clustering of circulating tumor cells confers high metastatic potential ^21,49,50^, and cell adhesion molecules K14, plakoglobin and E-cadherin have been shown to be required for metastasis ^15,21,22^. Therefore, K19 may also be part of breast cancer cell molecular machinery involved in maintaining CTC clusters during metastasis. As our data from migration, mammosphere and colony formation assays show, K19 seems to either promote or inhibit metastasis in a stage-specific manner.

The fact that K19 was required to form colonies in low attachment conditions may help explain interesting observations that have been made previously in regards to tumor metastasis. While the basal-like subtype of breast cancer is considered to be more invasive and aggressive than the luminal subtype ^43,51^, luminal-like cells without basal-like traits were fully capable of initiating invasive tumors in immune-deficient mice ^52^. In fact, phenotypically pure luminal-like breast cancer cells formed larger and more invasive tumors than basal-like cells ^52^. Also, increased levels of *KRT19* mRNA in CTCs during metastasis is associated with worse patient survival ^53–56^, and K19 has been shown to be even released by breast cancer cells of high metastatic potential ^57^. Still, as the lack of K19 expression is also correlated with worse survival of young women with triple-negative breast cancer ^58^, additional studies are needed to reconcile these seemingly conflicting observations. Differences in the role of K19 in tumor metastasis are likely due to the context-dependent role of K19 as breast cancer is a heterogeneous disease and metastasis itself is a complex process involving multiple factors ^2,4,6^.

K19 is a stem cell marker in several tissue types, including breast tissue ^59–61^, but its role in cancer stem cells is unclear. As mammosphere formation in low attachment conditions has been linked to cancer stem cell activities due to resistance to anoikis, the requirement of K19 to form mammospheres suggests that K19 plays an active role in maintaining stem-like properties of breast cancer cells. Consistent with our observation, K19-negative mammary progenitor cells showed delayed tumor onset and displayed lower metastatic potential in xenograft assay than K19-positive progenitor cells ^62^. As a subpopulation of cancer cells exhibiting stem-like properties are considered to be critical for metastasis ^63^, detailing the role of K19 in cancer stem cells will be important to study in the future.

In the absence of K19, E-cadherin was found in endocytic compartments indicating that most E-cadherin was internalized in the absence of K19. Indeed, the use of dynamin inhibitor to inhibit internalization of cell surface proteins including E-cadherin strengthened cell-cell adhesion of *KRT19* KO cells, suggesting that internalization of adhesion molecule is a culprit for defective cell-cell adhesion. Still, the mechanism of how K19 affects localization of E-cadherin remains unclear. Without keratins, desmosomes are smaller and consistent with this, plakoglobin levels were dependent on K19. Since a reduction of desmosomes precedes the loss of adherens junctions in tumors ^11,64,65^, a loss of K19 may internalize E-cadherin by first deregulating desmosomes which would then trigger the disassembly of adherens junctions. However, plakoglobin is also a part of adherens junction and K19 interacts with β-catenin ^24^. Therefore, K19 may directly stabilize adherens junction components at the cell surface independent of its effect on desmosomes. Future studies will need to elucidate the exact mechanism of how K19 regulates E-cadherin localization.

E-cadherin has long been considered as a tumor suppressor as it is one key component maintaining the epithelial state ^19^. However, recent evidences suggest that E-cadherin may promote breast cancer metastasis. While loss of E-cadherin increased invasion, it reduced cell proliferation and survival, CTC number, and metastasis in various models of invasive ductal carcinomas ^66^. In addition, knockdown of E-cadherin abrogated mammosphere formation of MCF7 cells ^67^. Therefore, the parallel between K19 and E-cadherin in inhibition of cell migration and promotion of cell proliferation and mammosphere formation ^32,66^, together with regulation of E-cadherin localization by K19 suggest that K19 may functionally interact with E-cadherin to promote breast cancer metastasis.

## Conclusions

Our study demonstrates that K19 is required to maintain cell morphology and cell-cell adhesion in MCF7 breast cancer cells. Cells lacking K19 show defects in plakoglobin expression, E-cadherin-β-catenin interaction and E-cadherin localization. Importantly, while K19 inhibited cell migration, it was required to form colonies in low adherent conditions. These data suggest that regulation of cell-cell mechanics of cells by K19 differentially affects processes critical to cancer metastasis.

## Materials and Methods

### Cell lines used

MCF7 *KRT19* KO cells generated using the CRISPR/Cas9 system and MCF7 *KRT19* KO cells stably expressing pLenti CMV/TO hygro empty vector or *KRT19* have been described previously ^32^. For MCF7 *KRT19* KO cells stably expressing GFP or GFP-K19, human *KRT19* was cloned out of pMRB101 plasmid (courtesy of Dr. Bishr Omary, Rutgers University) and cloned into pLenti CMV/TO hygro (Addgene, Cambridge, MA) using an In-Fusion HD cloning system (Takara, Mountain View, CA) with oligonucleotides: 5’-TCAGTCGACT<u>GGATCC</u>ATGGTGAGCAAGGGCGAGG-3’ (BamHI site underlined) and 5’-GAAAGCTGGG<u>TCTAG</u>TCAGAGGACCTTGGAGGCAG-3’ (XbaI site underlined), following the manufacturer’s protocol.

### Lentiviral supernatants

Lentiviral supernatants were generated as described previously ^32^. Lentiviral supernatants, collected 24 h after transfection, were used to infect subconfluent MCF7 *KRT19* KO cells (KO2) in three sequential 4 h incubation in the presence of 4 µg/ml polybrene (Sigma-Aldrich, St Louis, MO). Transductants were selected in hygromycin (100 µg/ml), beginning 48 h after infection.

### Cell culture and cell lines used

Dulbecco’s Modified Essential Medium (VWR Life Science, Carlsbad, CA) medium was used for MCF7 (ATCC, Manassas VA) growth in 37 °C incubator supplied with 5 % CO_2_. Medium was supplemented with 10% fetalgro bovine growth serum (RMBIO, Missoula, MT), and 100 units/ml penicillin-100 μg/ml streptomycin (GE Healthcare, Logan UT). MCF7 cells used were authenticated to be 100% match to MCF7 cells (ATCC, HTB-22) by short-tandem repeat profiling service (date performed: 12/18/18). 100 µg/ml hygromycin was added to medium of cells stably expressing pLenti CMV/TO Hygro plasmids.

### Antibodies

The following antibodies: anti-K19 (A-53), anti-K18 (C-04), anti-GAPDH (FL-335), anti-β catenin (15B8), anti-E-cadherin (67A4), anti-β-actin (C4), anti-LAMP-1(H4A3), anti-mouse IgG, and anti-rabbit IgG were from Santa Cruz Biotechnology (Santa Cruz, CA); anti-δ-1 catenin (D7S2M) and anti-plakoglobin (D9M1Q) were from Cell Signaling Technology (Danvers, MA); anti-α-tubulin was from Proteintech (Rosemont, IL); anti-GFP (12A6) was from the Developmental Studies Hybridoma Bank (Iowa City, IA).

### Cells shape and adhesion assessment

15,000 cells were plated on each well in a 6-well plate. 24 h after plating, at least five random fields of cells in culture were taken using a phase contrast microscope (Olympus optical company. LTD, Japan). To assess cell shape, cells were categorized as either round or elongated. For cell adhesion, cells were categorized into three types (high, medium, or low) based on their attachments to surrounding cells. To quantify number of cells per cluster, cells in contact with adjoining cells were counted a day after passaging. Total of five experiments were analyzed.

### Digital holographic microscope (DHM)

30,000 cells in were plated in each of 35 mm glass bottom plates. The next day, images were taken using DHM as described previously ^68^. The DHM system utilized a wavelength 633 mm HeNe laser to generate reference and objective wave beams and created a lateral resolution of 1.2 µm with a pixel scale of 0.18 µm/pixel. The holograms were captured by a 1.3 MP CMOS camera (Lumenera Corporation, Inc., Ontario, Canada). Detailed information about the set up was published in ^45,68,69^. Cell phase maps of parental and *KRT19* KO2 cells cultured on glass were collected with a sample size of n = 259 and 173, respectively. Phase images of each individual cell were segmented and 17 phase parameters were extracted (Supplementary Table 1) using an in-house MATLAB code published in ^70^. Representative cells were selected by measuring the sum of differences of all parameters of each individual cell to the population’s mean, and selecting the cells with the lowest measured values ^68^.

### Dispase assay

At 100% confluency, cells in a 6 well plate were washed with 1X PBS then subjected to 2.5 units/ml of dispase (Stemcell Technologies, Kent, WA). The plate was placed on an orbital shaker to induce mechanical stress at room temperature. After 40 min, number of fragments was counted by naked eyes. Images of the plate were taken using ChemiDoc Touch Imager (Bio-Rad, Hercules, CA). Each experiment contained at least three replicates per condition, and every experiment was performed at least three times. For dynasore (Ambeed, Inc, Arlington Heights, IL) treatment, 100 µM of dynasore was added for 2 h in each well before the dispase treatment.

### Preparation of cell lysates, protein gel electrophoresis, and immunoblotting

Cells grown on tissue culture plates were washed with PBS and prepared in cold Triton lysis buffer (1 % Triton X-100, 40 mM HEPES (pH 7.5), 120 mM sodium chloride, 1 mM EDTA, 1 mM phenyl methylsulfonyl fluoride, 10 mM sodium pyrophosphate, 1 μg/ml each of cymostatin, leupeptin and pepstatin, 10 μg/ml each of aprotinin and benzamidine, 2 μg/ml antipain, 1 mM sodium orthovanadate, 50 mM sodium fluoride). For immunoblotting, cell lysates were centrifuged to remove cell debris. Protein concentration was determined using the Bio-Rad Protein Assay (Bio-Rad) with BSA as standard then were prepared in Laemmli SDS-PAGE sample buffer. Aliquots of protein lysate were resolved by SDS-PAGE, transferred to nitrocellulose membranes (0.45 μm) (Bio-Rad, Hercules, CA) and immunoblotted with the indicated antibodies, followed by horseradish peroxidase-conjugated goat anti-mouse or goat anti-rabbit IgG (Sigma-Aldrich) and Amersham ECL Select Western Blotting Detection Reagent or Pierce ECL Western Blotting Substrate (Thermo Fisher Scientific, Hudson, NH). Signals were detected using ChemiDoc Touch Imager (Bio-Rad) or CL1500 Imaging System (Thermo Fisher Scientific). For Western blot signal quantitation, the Image Lab software (Bio-Rad) was used.

### Immunoprecipitation

Cells lysates were prepared using 1 % triton lysis buffer and precleared with prewashed Protein G Sepharose (PGS) beads (GE Healthcare) for 30 min on a shaker at 4 °C. Then, prewashed lysates were incubated with an indicated antibody on a shaker for 4 h in 4 °C. Samples were then incubated for 45 min with prewashed PGS beads. Afterwards, beads were spun down, washed and prepared for protein gel electrophoresis and immunoblotting.

### Biotin labeling of cell surface proteins

Cells were washed with ice-cold 20 mM HEPES, pH 7.5 in 1X PBS then treated with 400 μg/ml sulfo-*N*-hydroxysulfosuccinimide-biotin (Thermo Fisher Scientific) prepared in the washing buffer for 40 min on an orbital shaker at 4°C. A duplicate set of cells was kept without biotin as a negative control. Cells were then lysed with prechilled 1 % triton lysis buffer. Immunoprecipitation was performed with prewashed neutravidin agarose beads (Thermo Fisher Scientific) for 1 h at 4 °C. Samples were then prepared for protein gel electrophoresis and immunoblotting.

### Immunofluorescence (IF) staining

Cells grown on glass coverslips (VWR) were washed in PBS, fixed in 4% paraformaldehyde in PBS, and permeabilized in 0.1% Triton X-100. Samples were blocked in 5% normal goat serum (NGS, RMBIO) in PBS before staining with primary antibodies diluted in blocking buffer and a mixture of 1:1000 of Alexa Fluor 488-conjugated goat anti-mouse secondary antibody (Invitrogen), 1:5000 Dapi (Sigma-Aldrich), and/or 1:400 Phalloidin (MilliporeSigma, Burlington, MA) in 1X PBS was added for 1 h incubation at RT. After 1X PBS washes, coverslips were mounted on microscope slides with mounting medium containing 1,4-diaza-bicyclo[2.2.2]octane (Electron Microscopy Sciences, Hatfield, PA). Fluorescence images were taken using the Olympus optical elements fluorescence microscope (Olympus). For transferrin labelling, 10 μg/ml of Alexa Fluor 594-conjugated transferrin was loaded to cells for 30 min prior to fixation. For E-cadherin and LAMP1 double staining, cells immunostained for E-cadherin with Alexa Fluor 488 secondary antibody were blocked with 26 µg/ml of AffiniPure Fab Fragment Goat Anti-Mouse IgG (H+L) (Jackson ImmunoResearch, Inc, West Grove, PA) in 5% NGS in PBS overnight before being stained for LAMP1 with Alexa Fluor 594-conjugated goat anti-mouse secondary antibody (Invitrogen).

### Calcium or serum depletion and re-stimulation

15,000 cells were plated on each well in a 12 wells plate. Next day, cells were placed in calcium free, low glucose with L-glutamine DMEM (USBiological life science, Salem, MA) or 0.1 % serum-containing medium for time indicated in each figure. Cells were then either left unstimulated or stimulated with 5 mM of CaCl_2_ or 10 % serum for indicated time in figures. Images of either live cells in culture or cells stained with crystal violet were taken using light microscope (Olympus). For crystal violet staining of cells, a mixture of 0.1 % of crystal violet and 10 % ethanol prepared in 1X PBS was added to cells, which were then placed on a shaker at room temperature for 15 min. Cells were washed with 1X PBS before imaging.

### Wound healing assay

Wounds were made on cells grown to a confluent monolayer on each of a 6 well plate using a pipet tip. Cells were washed with the medium and images were taken using the light microscope (Olympus) at different time points until wounds were closed. Percentage of wound closure over time was calculated by measuring wound area using a free hand tool of the ImageJ software and normalizing area to the area of the initial wound. Each experiment contained at least three replicates per condition, and every experiment was performed at least three times.

### Mammosphere formation assay

100,000 cells were plated on each well of an ultra-low attachment plate (Corning Life Sciences, Corning, NY). After 7 days, images were taken using the light microscope (Olympus). Total area of mammospheres were obtained using the ImageJ software. Each experiment contained at least three replicates per condition, and every experiment was performed at least three times.

### Anchorage-independent growth in soft agar

Soft agar colony growth assay was performed as previously described ^71^. 20,000 cells were plated per well in a 6-well plate in 1 ml of 0.1% agarose in media on top of a 2-ml bottom layer of 0.5% agarose in media. Plates were incubated at 37 °C in incubator for one week. Cells were fed every other day and images were taken using ChemiDoc Touch Imager (Bio-Rad) or CL1500 Imaging System (Thermo Fisher Scientific). Colony area was measured using the ImageJ software. Each experiment contained at least three replicates per condition, and every experiment was performed at least three times.

### MCF7 RNA-sequencing and bioinformatic analyses

RNA-sequencing of parental and *KRT19* KO MCF7 cells was performed previously ^32^. List of genes was sorted based on q-value (less than 0.05), and fold change (greater than 1.5). 387 genes upregulated in *KRT19* KO cells were uploaded in functional annotation tool (DAVID software) and keywords in functional categories was selected. Top thirteen pathways were sorted based on gene number.

### Graphs and statistics

Mean ± standard error of means was used. For comparisons between two conditions, Student’s t-test was performed to test the statistical significance.

## Author Contributions

Conceptualization, B.M.C; methodology, S.A., V.L., T.P., G.N., C.B.R., and B.M.C.; validation, S.A., P.S., and K.B.; formal analysis, S.A., P.S., K.B., R.M., M.C., A.F., T.P., P.D., C.B.R., and B.M.C.; investigation, S.A., P.S., K.B., and T.P.; resources, G.N., C.B.R., and B.M.C.; data curation, S.A., P.S., K.B., V.L., C.B.R., and B.M.C.; writing-original draft preparation, S.A. and B.M.C.; writing-review and editing, S.A., C.B.R., and B.M.C.; visualization, S.A. and B.M.C.; supervision, B.M.C; project administration, B.M.C.; funding acquisition, C.B.R. and B.M.C.

## Funding

This work was supported by National Cancer Institute under grant number R15CA2113071 (to B.M.C.) and National Institute of Biomedical Imaging and Bioengineering under grant number R03EB028017 (to C.R.).

## Acknowledgments

We thank the Chung lab members for their support and Dr. Pamela Tuma of the Department of Biology at The Catholic University of America for sharing instruments.

## Conflicts of Interest

The authors declare no conflict of interest.

## Notes

### Competing Interest Statement

The authors have declared no competing interest.

## References

1. Hanahan D, Weinberg RA. The hallmarks of cancer. Cell 2000; 100:57–70.

2. Gupta GP, Massagué J. Cancer Metastasis: Building a Framework. Cell 2006; 127:679–95.

3. Dillekås H, Rogers MS, Straume O. Are 90% of deaths from cancer caused by metastases? Cancer Med 2019; 8:5574–6.

4. Lambert AW, Pattabiraman DR, Weinberg RA. Emerging Biological Principles of Metastasis. Cell 2017; 168:670–91.

5. Sharma P, Alsharif S, Fallatah A, Chung BM. Intermediate Filaments as Effectors of Cancer Development and Metastasis: A Focus on Keratins, Vimentin, and Nestin. Cells 2019; 8.

6. Weigelt B, Peterse JL, van’t Veer LJ. Breast cancer metastasis: markers and models. NatRevCancer 2005; 5:591–602.

7. Desai BV, Harmon RM, Green KJ. Desmosomes at a glance. JCellSci 2009; 122:4401–7.

8. Green KJ, Jaiganesh A, Broussard JA. Desmosomes: Essential contributors to an integrated intercellular junction network. F1000Res 2019; 8.

9. Seltmann K, Fritsch AW, Kas JA, Magin TM. Keratins significantly contribute to cell stiffness and impact invasive behavior. ProcNatlAcadSciUSA 2013; 110:18507–12.

10. Huang RY-J, Guilford P, Thiery JP. Early events in cell adhesion and polarity during epithelial-mesenchymal transition. J Cell Sci 2012; 125:4417–22.

11. Kröger C, Loschke F, Schwarz N, Windoffer R, Leube RE, Magin TM. Keratins control intercellular adhesion involving PKC-α–mediated desmoplakin phosphorylation. J Cell Biol 2013; 201:681–92.

12. Chung BM, Rotty JD, Coulombe PA. Networking galore: intermediate filaments and cell migration. CurrOpinCell Biol 2013; 25:600–12.

13. Leduc C, Etienne-Manneville S. Intermediate filaments in cell migration and invasion: the unusual suspects. CurrOpinCell Biol 2015; 32:102–12.

14. Wang F, Chen S, Liu HB, Parent CA, Coulombe PA. Keratin 6 regulates collective keratinocyte migration by altering cell-cell and cell-matrix adhesion. J Cell Biol 2018;

15. Cheung KJ, Gabrielson E, Werb Z, Ewald AJ. Collective invasion in breast cancer requires a conserved basal epithelial program. Cell 2013; 155:1639–51.

16. Chung BM, Arutyunov A, Ilagan E, Yao N, Wills-Karp M, Coulombe PA. Regulation of C-X-C chemokine gene expression by keratin 17 and hnRNP K in skin tumor keratinocytes. JCell Biol 2015; 208:613–27.

17. Cheung KJ, Ewald AJ. Illuminating breast cancer invasion: diverse roles for cell-cell interactions. CurrOpinCell Biol 2014; 30:99–111.

18. Yang Y, Zheng H, Zhan Y, Fan S. An emerging tumor invasion mechanism about the collective cell migration. Am J Transl Res 2019; 11:5301–12.

19. Tsai JH, Yang J. Epithelial–mesenchymal plasticity in carcinoma metastasis. Genes Dev 2013; 27:2192–206.

20. Plaks V, Koopman CD, Werb Z. Circulating Tumor Cells. Science [Internet] 2013 [cited 2020 Mar 15]; 341. Available from: https://www.ncbi.nlm.nih.gov/pmc/articles/PMC3842225/

21. Aceto N, Bardia A, Miyamoto DT, Donaldson MC, Wittner BS, Spencer JA, Yu M, Pely A, Engstrom A, Zhu H, et al. Circulating tumor cell clusters are oligoclonal precursors of breast cancer metastasis. Cell 2014; 158:1110–22.

22. Padmanaban V, Krol I, Suhail Y, Szczerba BM, Aceto N, Bader JS, Ewald AJ. E-cadherin is required for metastasis in multiple models of breast cancer. Nature 2019; 573:439–44.

23. Bambang IF, Lu D, Li H, Chiu L-L, Lau QC, Koay E, Zhang D. Cytokeratin 19 regulates endoplasmic reticulum stress and inhibits ERp29 expression via p38 MAPK/XBP-1 signaling in breast cancer cells. Exp Cell Res 2009; 315:1964–74.

24. Saha SK, Choi HY, Kim BW, Dayem AA, Yang G-M, Kim KS, Yin YF, Cho S-G. KRT19 directly interacts with β-catenin/RAC1 complex to regulate NUMB-dependent NOTCH signaling pathway and breast cancer properties. Oncogene 2017; 36:332–49.

25. Ju J-H, Yang W, Lee K-M, Oh S, Nam K, Shim S, Shin SY, Gye MC, Chu I-S, Shin I. Regulation of cell proliferation and migration by keratin19-induced nuclear import of early growth response-1 in breast cancer cells. Clin Cancer Res 2013; 19:4335–46.

26. Bhagirath D, Zhao X, West WW, Qiu F, Band H, Band V. Cell type of origin as well as genetic alterations contribute to breast cancer phenotypes. Oncotarget 2015; 6:9018–30.

27. Dai X, Cheng H, Bai Z, Li J. Breast Cancer Cell Line Classification and Its Relevance with Breast Tumor Subtyping. J Cancer 2017; 8:3131–41.

28. Moll R, Franke WW, Schiller DL, Geiger B, Krepler R. The catalog of human cytokeratins: Patterns of expression in normal epithelia, tumors and cultured cells. Cell 1982; 31:11–24.

29. Comsa S, Cîmpean AM, Raica M. The Story of MCF-7 Breast Cancer Cell Line: 40 years of Experience in Research. Anticancer Res 2015; 35:3147–54.

30. Cowin P, Kapprell HP, Franke WW. The complement of desmosomal plaque proteins in different cell types. The Journal of Cell Biology 1985; 101:1442–54.

31. Gloushankova NA, Rubtsova SN, Zhitnyak IY. Cadherin-mediated cell-cell interactions in normal and cancer cells. Tissue Barriers 2017; 5:e1356900.

32. Sharma P, Alsharif S, Bursch K, Parvathaneni S, Anastasakis DG, Chahine J, Fallatah A, Nicolas K, Sharma S, Hafner M, et al. Keratin 19 regulates cell cycle pathway and sensitivity of breast cancer cells to CDK inhibitors. Sci Rep 2019; 9:14650.

33. Lam VK, Nguyen TC, Chung BM, Nehmetallah G, Raub CB. Quantitative assessment of cancer cell morphology and motility using telecentric digital holographic microscopy and machine learning. Cytometry A 2018; 93:334–45.

34. Lam VK, Nguyen T, Phan T, Chung B-M, Nehmetallah G, Raub CB. Machine Learning with Optical Phase Signatures for Phenotypic Profiling of Cell Lines. Cytometry Part A [Internet] 2019 [cited 2019 May 9]; Available from: https://onlinelibrary.wiley.com/doi/abs/10.1002/cyto.a.23774

35. Pece S, Gutkind JS. E-cadherin and Hakai: signalling, remodeling or destruction? Nat Cell Biol 2002; 4:E72–4.

36. Le TL, Yap AS, Stow JL. Recycling of E-Cadherin: A Potential Mechanism for Regulating Cadherin Dynamics. The Journal of Cell Biology 1999; 146:219–32.

37. Paterson AD, Parton RG, Ferguson C, Stow JL, Yap AS. Characterization of E-cadherin Endocytosis in Isolated MCF-7 and Chinese Hamster Ovary Cells THE INITIAL FATE OF UNBOUND E-CADHERIN. J Biol Chem 2003; 278:21050–7.

38. Ponti D, Costa A, Zaffaroni N, Pratesi G, Petrangolini G, Coradini D, Pilotti S, Pierotti MA, Daidone MG. Isolation and in vitro propagation of tumorigenic breast cancer cells with stem/progenitor cell properties. Cancer Res 2005; 65:5506–11.

39. Moll R, Divo M, Langbein L. The human keratins: biology and pathology. HistochemCell Biol 2008; 129:705–33.

40. Eswaran J, Cyanam D, Mudvari P, Reddy SD, Pakala SB, Nair SS, Florea L, Fuqua SA, Godbole S, Kumar R. Transcriptomic landscape of breast cancers through mRNA sequencing. SciRep 2012; 2:264.

41. Lehmann BD, Bauer JA, Chen X, Sanders ME, Chakravarthy AB, Shyr Y, Pietenpol JA. Identification of human triple-negative breast cancer subtypes and preclinical models for selection of targeted therapies. The Journal of clinical investigation 2011; 121:2750–67.

42. Perou CM, Sørlie T, Eisen MB, Rijn M van de, Jeffrey SS, Rees CA, Pollack JR, Ross DT, Johnsen H, Akslen LA, et al. Molecular portraits of human breast tumours. Nature 2000; 406:747–52.

43. Sorlie T, Perou CM, Tibshirani R, Aas T, Geisler S, Johnsen H, Hastie T, Eisen MB, van de Rijn M, Jeffrey SS, et al. Gene expression patterns of breast carcinomas distinguish tumor subclasses with clinical implications. ProcNatlAcadSciUSA 2001; 98:10869–74.

44. Bhagirath D, Zhao X, Mirza S, West WW, Band H, Band V. Mutant PIK3CA Induces EMT in a Cell Type Specific Manner. PLoS ONE 2016; 11:e0167064.

45. Lam VK, Sharma P, Nguyen T, Nehmetallah G, Raub CB, Chung BM. Morphology, Motility, and Cytoskeletal Architecture of Breast Cancer Cells Depend on Keratin 19 and Substrate. Cytometry A 2020;

46. König K, Meder L, Kröger C, Diehl L, Florin A, Rommerscheidt-Fuss U, Kahl P, Wardelmann E, Magin TM, Buettner R, et al. Loss of the Keratin Cytoskeleton Is Not Sufficient to Induce Epithelial Mesenchymal Transition in a Novel KRAS Driven Sporadic Lung Cancer Mouse Model. PLOS ONE 2013; 8:e57996.

47. Mendez MG, Kojima S, Goldman RD. Vimentin induces changes in cell shape, motility, and adhesion during the epithelial to mesenchymal transition. FASEB J 2010; 24:1838–51.

48. Mori S, Chang JT, Andrechek ER, Matsumura N, Baba T, Yao G, Kim JW, Gatza M, Murphy S, Nevins JR. Anchorage-independent cell growth signature identifies tumors with metastatic potential. Oncogene 2009; 28:2796–805.

49. Giuliano M, Shaikh A, Lo HC, Arpino G, Placido SD, Zhang XH, Cristofanilli M, Schiff R, Trivedi MV. Perspective on Circulating Tumor Cell Clusters: Why It Takes a Village to Metastasize. Cancer Res 2018; 78:845–52.

50. Gkountela S, Castro-Giner F, Szczerba BM, Vetter M, Landin J, Scherrer R, Krol I, Scheidmann MC, Beisel C, Stirnimann CU, et al. Circulating Tumor Cell Clustering Shapes DNA Methylation to Enable Metastasis Seeding. Cell 2019; 176:98–112.e14.

51. Badve S, Dabbs DJ, Schnitt SJ, Baehner FL, Decker T, Eusebi V, Fox SB, Ichihara S, Jacquemier J, Lakhani SR, et al. Basal-like and triple-negative breast cancers: a critical review with an emphasis on the implications for pathologists and oncologists. Mod Pathol 2011; 24:157–67.

52. Kim J, Villadsen R, Sørlie T, Fogh L, Grønlund SZ, Fridriksdottir AJ, Kuhn I, Rank F, Wielenga VT, Solvang H, et al. Tumor initiating but differentiated luminal-like breast cancer cells are highly invasive in the absence of basal-like activity. PNAS 2012; 109:6124–9.

53. Kahn HJ, Yang L-Y, Blondal J, Lickley L, Holloway C, Hanna W, Narod S, McCready DR, Seth A, Marks A, et al. RT-PCR amplification of CK19 mRNA in the blood of breast cancer patients: correlation with established prognostic parameters. Breast Cancer Res Treat 2000; 60:143–51.

54. Yu X-F, Yang H-J, Lei L, Wang C, Huang J. CK19 mRNA in blood can predict non-sentinel lymph node metastasis in breast cancer. Oncotarget 2016; 7:30504–10.

55. Saloustros E, Perraki M, Apostolaki S, Kallergi G, Xyrafas A, Kalbakis K, Agelaki S, Kalykaki A, Georgoulias V, Mavroudis D. Cytokeratin-19 mRNA-positive circulating tumor cells during follow-up of patients with operable breast cancer: prognostic relevance for late relapse. Breast Cancer Res 2011; 13:R60.

56. Reinholz MM, Kitzmann KA, Tenner K, Hillman D, Dueck AC, Hobday TJ, Northfelt DW, Moreno-Aspitia A, Roy V, LaPlant B, et al. Cytokeratin-19 and Mammaglobin Gene Expression in Circulating Tumor Cells from Metastatic Breast Cancer Patients Enrolled in North Central Cancer Treatment Group Trials, N0234/336/436/437. Clin Cancer Res 2011; 17:7183–93.

57. Alix-Panabieres C, Vendrell JP, Slijper M, Pelle O, Barbotte E, Mercier G, Jacot W, Fabbro M, Pantel K. Full-length cytokeratin-19 is released by human tumor cells: a potential role in metastatic progression of breast cancer. Breast Cancer Res 2009; 11:R39.

58. Parikh RR, Yang Q, Higgins SA, Haffty BG. Outcomes in Young Women With Breast Cancer of Triple-Negative Phenotype: The Prognostic Significance of CK19 Expression. International Journal of Radiation Oncology • Biology • Physics 2008; 70:35–42.

59. Asfaha S, Hayakawa Y, Muley A, Stokes S, Graham TA, Ericksen RE, Westphalen CB, von Burstin J, Mastracci TL, Worthley DL, et al. Krt19(+)/Lgr5(-) Cells Are Radioresistant Cancer-Initiating Stem Cells in the Colon and Intestine. Cell stem cell 2015; 16:627–38.

60. Dittmer J, Rody A. Cancer stem cells in breast cancer. HistolHistopathol 2013; 28:827–38.

61. Petersen OW, Polyak K. Stem cells in the human breast. Cold Spring Harb PerspectBiol 2010; 2:a003160.

62. Bhagirath D, Zhao X, West WW, Qiu F, Band H, Band V. Cell type of origin as well as genetic alterations contribute to breast cancer phenotypes. Oncotarget 2015; 6:9018–30.

63. Baccelli I, Trumpp A. The evolving concept of cancer and metastasis stem cells. J Cell Biol 2012; 198:281–93.

64. Dusek RL, Attardi LD. Desmosomes: new perpetrators in tumour suppression. Nat Rev Cancer 2011; 11:317–23.

65. Wallace L, Roberts-Thompson L, Reichelt J. Deletion of K1/K10 does not impair epidermal stratification but affects desmosomal structure and nuclear integrity. Journal of Cell Science 2012; 125:1750–8.

66. Padmanaban V, Krol I, Suhail Y, Szczerba BM, Aceto N, Bader JS, Ewald AJ. E-cadherin is required for metastasis in multiple models of breast cancer. Nature 2019; 573:439–44.

67. Iglesias JM, Beloqui I, Garcia-Garcia F, Leis O, Vazquez-Martin A, Eguiara A, Cufi S, Pavon A, Menendez JA, Dopazo J, et al. Mammosphere Formation in Breast Carcinoma Cell Lines Depends upon Expression of E-cadherin. PLOS ONE 2013; 8:e77281.

68. Lam VK, Nguyen T, Phan T, Chung BM, Nehmetallah G, Raub CB. Machine Learning with Optical Phase Signatures for Phenotypic Profiling of Cell Lines. Cytometry Part A 2019; 95:757–68.

69. Nguyen T, Nehmetallah G, Raub C, Mathews S, Aylo R. Accurate quantitative phase digital holographic microscopy with single- and multiple-wavelength telecentric and nontelecentric configurations. Applied Optics 2016; 55:5666–83.

70. Lam VK, Nguyen TC, Chung BM, Nehmetallah G, Raub CB. Quantitative assessment of cancer cell morphology and motility using telecentric digital holographic microscopy and machine learning. Cytometry Part A 2018; 93:334–45.

71. Chung BM, Dimri M, George M, Reddi AL, Chen G, Band V, Band H. The role of cooperativity with Src in oncogenic transformation mediated by non-small cell lung cancer-associated EGF receptor mutants. Oncogene 2009; 28:1821–32.

