## Supplemental Data for "Keratin 19 maintains E-cadherin localization at the cell surface and stabilizes cell-cell adhesion of MCF7 cells"

**Supplementary Data**

| Parameters | Full name | parental | *KRT19* KO | p-value |
|---|---|---|---|---|
| PhaseMean* | Average phase height | 3950.0143 | 3638.2445 | 0.001 |
| PhaseMean SD | Phase height standard deviation | 1965.247 | 2032.7613 | 0.229 |
| ku | Phase height kurtosis | 2.2347166 | 2.2688439 | 0.649 |
| sk* | Phase height skewness | -0.0868383 | 0.1012174 | 0 |
| Area (um2)* | Segmented cell area | 460.37573 | 562.81516 | 0 |
| Perimeter (μm)* | Segmented cell perimeter | 95.032022 | 128.03663 | 0 |
| Eccentricity* | Segmented cell fit ellipse eccentricity | 0.7245655 | 0.8664618 | 0 |
| Form factor* | Segmented form factor | 0.65458 | 0.462963 | 0 |
| Contrast* | 2nd order texture parameter | 305.3581 | 329.5333 | 0.03 |
| Correlation | 2nd order texture parameter | 0.943321 | 0.9389649 | 0.07 |
| Energy | 2nd order texture parameter | 0.0001539 | 0.0001515 | 0.723 |
| Homogenity | 2nd order texture parameter | 0.1841623 | 0.1816254 | 0.364 |
| NuPhaseMx | Maximum nucleus phase height | 9107.7722 | 9529.5549 | 0.103 |
| NuPhaseAve | Average nucleus phase height | 5727.4355 | 5454.9353 | 0.062 |
| NuclearArea (um2)* | Segmented nucleus area | 185.03516 | 225.39083 | 0 |
| NuclearKu* | Phase height kurtosis | 0.614367 | 0.6625169 | 0.001 |
| NuclearSk* | Phase height skewness | 1.6379479 | 1.8013399 | 0.05 |

**Table S1. Cell parameters generated from DHM segmentation.** Circularity is determined by eccentricity and form factor. A perfect circle has eccentricity equal to 0 and form factor equal to 1. Means from three experimental repeats are shown. Differences are not statistically significant unless denoted by *$p < 0.05$; **$p < 0.001$

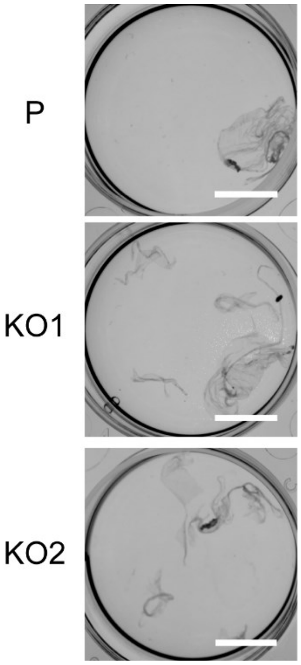


**Figure S1. *KRT19* KO cells show weakened cell-cell adhesion.** Phase contrast images of dispase treated cell fragments used for Fig 2E. Bar, 10 mm.

**A**

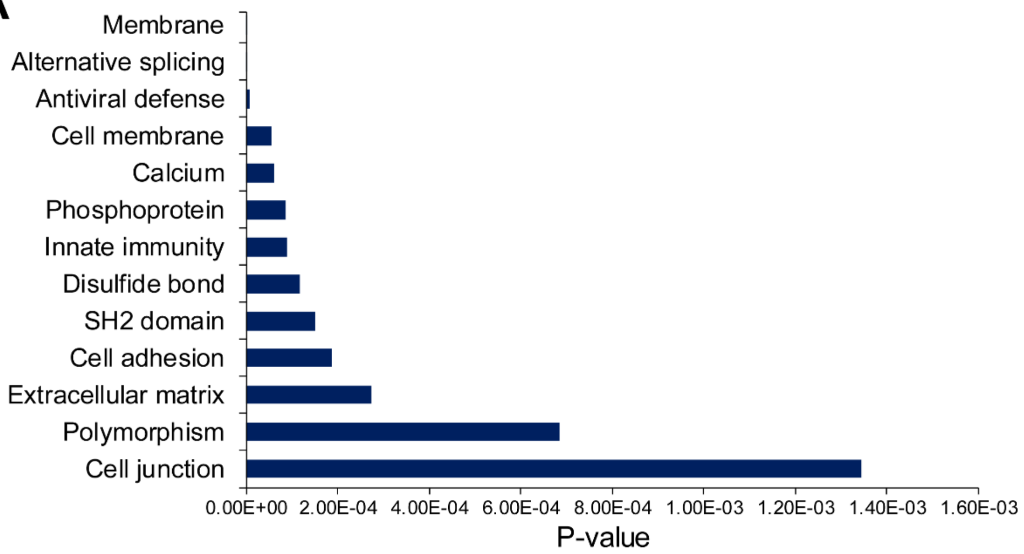


**B**

| Gene symbol | Gene name | RPKM P | RPKM KO | Log2 (fold change) |
|---|---|---|---|---|
| ENG | Endoglin | 10.5734 | 4.31752 | -1.292165 |
| ICAM1 | Intercellular Adhesion Molecule 1 | 1.60904 | 0.810933 | -0.988546 |
| RND3 | Rho Family GTPase 3 | 4.29229 | 2.21303 | -0.955725 |
| PCDH17 | Protocadherin 17 | 1.80447 | 0.945641 | -0.932211 |
| WISP2 | Cellular Communication Network Factor 5 | 16.1533 | 9.63394 | -0.745631 |
| JUP | Plakoglobin | 288.177 | 177.411 | -0.699860 |
| ITGAV | Integrin Subunit Alpha 5 | 9.5714 | 12.4971 | 0.384791 |
| PALLD | Palladin, cytoskeletal associated protein | 15.3946 | 20.5519 | 0.416847 |
| DSP | Desmoplakin | 54.1604 | 74.638 | 0.462672 |
| CLDN1 | Claudin1 | 4.56072 | 6.68239 | 0.551103 |
| CADM1 | Cell Adhesion Molecule 1 | 3.58198 | 5.59959 | 0.644564 |
| CDH3 | P-cadherin | 26.0338 | 42.2293 | 0.697858 |
| ITGA6 | Integrin Subunit Alpha 6 | 1.56782 | 2.54423 | 0.698469 |
| FSCN1 | Fascin Actin-bundling Protein 1 | 17.3017 | 28.4115 | 0.715561 |
| PODXL | Podocalyxin Like | 3.07798 | 5.14708 | 0.741770 |
| COL18A1 | Collagen Type XVIII Alpha 1 Chain | 14.3142 | 26.2243 | 0.873457 |
| CLDN9 | Claudin9 | 2.43677 | 4.46757 | 0.874520 |
| COL6A1 | Collagen Type VI Alpha 1 Chain | 4.91318 | 10.14 | 1.045329 |
| ADAM9 | ADAM Metallopeptidase Domain 9 | 9.97818 | 20.7437 | 1.055825 |
| DNMBP | Dynamin Binding Protein | 1.06723 | 2.31561 | 1.117521 |
| ATP1B1 | ATPase Na+/K+ Transporting Subunit Beta 1 | 7.76789 | 17.2509 | 1.151077 |
| MCAM | Melanoma Cell Adhesion Molecule | 1.89175 | 4.48657 | 1.245891 |
| CXCL12 | C-X-C Motif Chemokine Ligand 12 | 39.1316 | 97.7939 | 1.321410 |
| GPNMB | Glycoprotein Nmb | 1.16355 | 3.15029 | 1.436951 |
| COL5A1 | Collagen Type V Alpha 1 Chain | 2.04502 | 7.10348 | 1.796411 |
| MSLN | Pre-Pro-Megakaryocyte-Potentiating Factor | 0.9121 | 4.1815 | 2.196752 |
| TGFB2 | Transforming Growth Factor Beta 2 | 0.43267 | 3.88347 | 3.166021 |

**Figure S2. Increased expression of cell adhesion molecules in *KRT19* KO cells.** (**A**) A list of top keywords in functional categories associated with genes upregulated in *KRT19* KO from the RNA sequencing data [32] using the DAVID functional annotation software. (**B**) List of major cell adhesion molecules that are differentially regulated in *KRT19* KO cells.

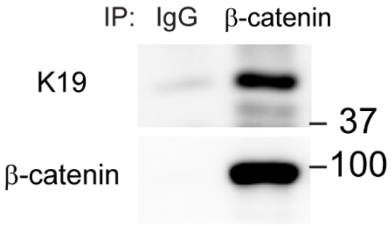


**Figure S3. K19 interacts with β-catenin in MCF7 cells.** IP with anti-β-catenin antibody or IgG control was performed. IP samples were subjected to SDS-PAGE and immunoblotting was performed with antibodies against K19 and β-catenin.

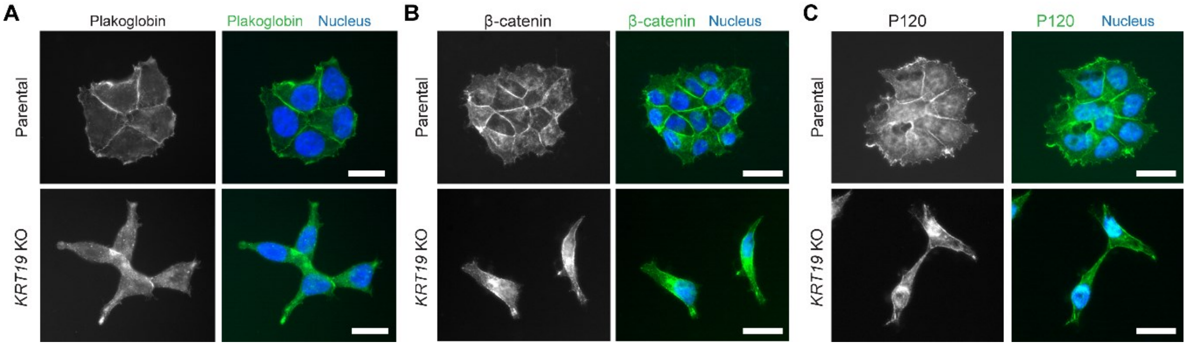


**Figure S4. Decreased membrane-bound plakoglobin, β-catenin, and P120-catenin in *KRT19* KO Cells.** Immunostaining of (**A**) plakoglobin, (**B**) β-catenin, and (**C**) P120-catenin in parental and *KRT19* KO (KO2) cells. Nuclei are shown in blue. Bar, 20 μm.

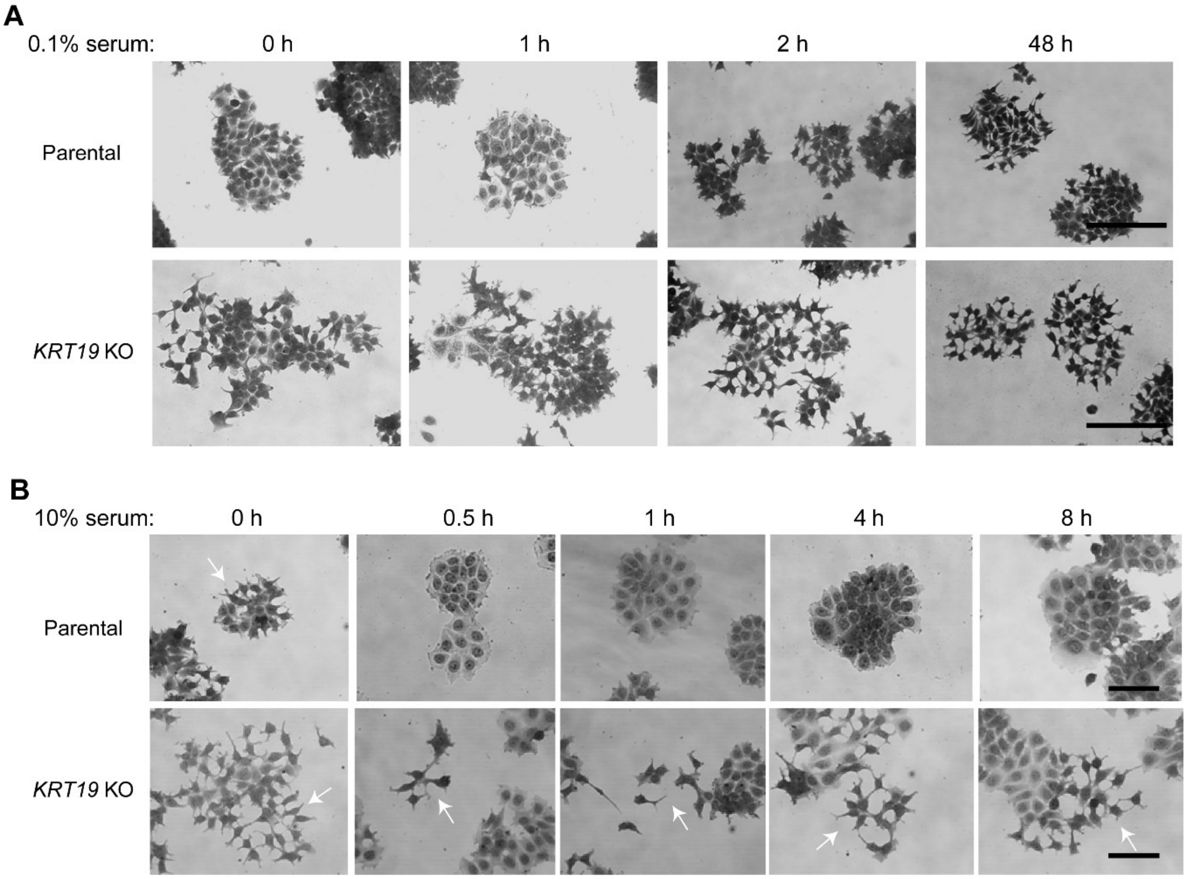


**Figure S5. Decreased responsiveness of *KRT19* KO cells for cell-cell adhesion upon serum stimulation.** (**A**) Phase contrast images of parental and *KRT19* KO (KO2) cells following serum starvation for indicated time points. Bar, 200 µm. Whereas parental cells show detachments between cells only after 48 h from serum starvation, shape of *KRT19* KO cells remain detached throughout serum starvation. Bar, 200 µm. (**B**) Phase contrast images of parental and *KRT19* KO (KO2) cells serum-starved for 24 h, then stimulated with 10% serum for indicated time points. Parental cells rapidly re-attach from 0.5 h time point but *KRT19* KO cells remain detached throughout serum starvation. Arrows indicate low cell-cell adhesions Bar, 100 µm.
